# Spatiotemporal Analysis of Category and Target-related Information Processing in the Brain during Object Detection

**DOI:** 10.1101/361642

**Authors:** Hamid Karimi-Rouzbahani, Ehsan Vahab, Reza Ebrahimpour, Mohammad Bagher Menhaj

## Abstract

To recognize a target object, the brain implements strategies which involve a combination of externally sensory-driven and internally task-driven mechanisms. While several studies have suggested a role for frontal brain areas in enhancing task-related representations in visual cortices, especially the lateral-occipital cortex, they remained silent about the type of information transferred to visual areas. However, the recently developed method of representational causality analysis, allowed us to track the movement of different types of information in the brain. Accordingly, we designed an EEG object detection experiment and evaluated the spatiotemporal dynamics of category- and target-related information across the brain using. Results showed that the prefrontal area initiated the processing of target-related information. This information was then transferred to posterior brain areas during stimulus presentation to facilitate object detection and to direct the decision-making procedure. We also observed that, as compared to category-related information, the target-related information could predict the behavioral detection performance more accurately, suggesting the dominant representation of internal compared to external information in brain signals. These results provided new evidence about the role of prefrontal cortices in the processing of task-related information the brain during object detection.

## Introduction

In order to recognize an object and initiate the relevant motor action, the brain must process a combination of the information provided by the external stimulus and the internal task goals. While a majority of classical studies have considered ‘object recognition’ as a feed-forward brain process (DiCarlo et al., 2012; VanRullen, 2012), recent studies have argued against this view by providing evidence about the significant influence of top-down task-dependent processes on bottom-up sensory-driven inputs (Harel et al., 2011; Bar et al., 2001; Bar et al., 2006; Vaziri-Pashkam and Xu, 2017). The effects of task are generally imposed on sensory processing by mechanisms involved in prediction (Summerfield et al., 2006), expectation (Puri et al., 2009; Meijs et al., 2018; Manahova et al., 2018) and attention (Stokes et al., 2009; Spyropoulos et al., 2018; Rohenkohl and Nobre, 2011; Battistoni et al., 2017; Summerfield and Egner, 2009; Paneri and Gregoriou, 2017).

Many previous studies have proposed that the visual information of an object is extracted in the brain by the transformations implemented in the ventral (from V1 area downward into the temporal lobe) as well as dorsal (from V1 area forward into the parietal lobe) visual streams (DiCarlo et al., 2012; Karimi-Rouzbahani et al., 2016; Karimi-Rouzbahani et al., 2017b). The abstracted input information is then sent to the prefrontal cortex, where the final decisions are made on the category of the perceived object or on other task-related aspects of information (eccentricity, distance, etc.; Hong et al., 2016; Kravitz et al., 2010; Schwarzlose et al., 2008). However, the task-related information, which is suggested to be mainly processed by the frontoparietal network, can drastically modulate object representations at many levels of the two streams, towards the subject’s goals (Woolgar et al., 2015a; Woolgar et al., 2015b; Jackson et al., 2017).

In object detection, which refers to the recognition of a target/cued object among a set of simultaneously or sequentially presented objects, the target-related information has been previously shown to exert effects on object-selective cortical areas such as the lateral occipital cortex (LOC; Kaiser et al., 2016; Soon et al., 2013; Stokes et al., 2009) and anterior inferior temporal cortex (Bansal et al., 2014), prior to (Soon et al., 2013; Battistoni et al., 2017; Milton and Pleydell-Pearce, 2016) and during (Kaiser et al., 2016; Goddard et al., 2016) the presentation of the object. A magnetoencephalography (MEG) study showed that the presence of target object in a scene could be significantly decoded as early as 160 ms post-stimulus whereas the information about non-target stimuli could not be decoded until after 200 ms post-stimulus (Kaiser et al., 2016). The target/non-target information was mainly localized to the LOC and authors suggested that the effect had been probably imposed by top-down signals from higher brain areas. However, this suggestion lacked quantitative support. Local filed potentials recorded from human temporal, parietal and frontal cortical areas during an object detection task, showed that the neural signals of object-selective populations, especially in inferior temporal cortex and fusiform gyri, could become significantly modulated after 250 ms post-stimulus (Bansal et al., 2014). This effect was most dominantly observed in the gamma band of neural activities, which had been previously suggested to provide the substrate for long-ranged transferring of information in the brain (Gregoriou et al., 2009). Accordingly, the authors associated the observed modulation to top-down signals from higher brain areas.

In order to gain a deeper insight into the spatiotemporal dynamics of object and task processing, a recent study, which fused MEG with functional magnetic resonance imaging (fMRI), found that a sequence of overlapping and separate structures along the frontoparietal and occipitotemporal cortices undertook task processing (Hebart et al., 2018). This observation was on par with previous findings which suggested that task processing was distributed along both the dorsal and ventral visual streams (Vaziri-Pashkam and Xu, 2017). Hebart et al. (2018) observed a late effect of task when decoded the brain signals with ignorable influence of task on object representations. They concluded that the task effects dominated object representations in downstream stages of the visual pathways. Yet, the authors did not study the possible causal role of frontal task-related information on visual areas.

Therefore, despite these recent endeavors, the spatiotemporal dynamics of target-related information processing across frontal and visual brain areas remains relatively ambiguous. Specifically, although these studies suggested the contribution of frontal brain areas to task-related effects found in visual areas, it is unknown if the frontal areas initiate the processing of target-related information first and whether (if at all) their signals are explicitly sent to sensory-processing areas. While a few studies have investigated the interactions between peri-frontal and peri-occipital brain areas in object/face/place recognition (Bar et al., 2006; Summerfield et al., 2006; Gregoriou et al., 2009), they overlooked to evaluate the nature of the signals which were transferred across the mentioned areas. More specifically, simultaneous recording from the frontal eye field (FEF) and area V4 of monkeys showed that attending to a stimulus within the shared receptive field of the recording sites could enhance the oscillatory coupling between the two areas, especially in the gamma frequency band (Gregoriou et al., 2009). The coupling was shown to be initiated by the FEF and affect activities in area V4 with around 8 to 13 ms latency. Therefore, the gamma band synchronization was suggested to provide a basis for communications across the two long-ranged areas during attentional deployment. Another study, using fMRI in humans observed a neural representation of the predicted perception in the medial frontal cortex, during a face-house discrimination task (Summerfield et al., 2006). More specifically, the authors observed an enhanced top-down connectivity between the frontal and face-selective visual areas when the subjects decided on the perception of faces, which supported a top-down predictive code for the detection of faces. An MEG-fMRI study, which investigated the spatiotemporal dynamics of object recognition in humans, suggested a two-pathway parallel model for object recognition in humans (Bar et al., 2006). In the model, the faster pathway carried low-spatial-frequency object information from subcortical/early visual cortex to orbitofrontal cortex and then to back to fusiform gyri. The slower pathway, on the other hand, which was the ventral visual stream, carried high-frequency object information. The information from the two pathways were combined at the fusiform gyri to provide a unified perception. The connectivity between the orbitofrontal and visual cortices was investigated based on the shifting power of the signals.

As it is obvious, the mentioned studies evaluated the connectivity between the frontal and the visual areas based on some implicit measures such as gamma-band synchronization (Gregoriou et al., 2009), shifting power (Bar et al., 2006) or causality in the activity signals (Summerfield et al., 2006), but not the information contained in the transferred signals. Therefore, we have remained uncertain about the task-relevance of these connectivities. Moreover, as these studies concentrated on some specific areas (Gregoriou et al., 2009) or suffered from low temporal resolution of fMRI (Summerfield et al., 2006), we could not evaluate the effects on the whole-brain scale or assess the temporal dynamics of information flow across different brain areas, respectively.

Fortunately, however, the recently-developed causality-based method of multi-variate representational similarity analysis (RSA; Goddard et al., 2016), when combined with the temporally high-resolution electroencephalography (EEG) and MEG, can allow the evaluation of target-related information flow across different brain regions. As a support for the performance of this method, a recent study, which first proposed the method, used it to reappraise the results reported by Bar et al. (2006) about the above-mentioned parallel visual processing pathways, and observed a totally different temporal dynamics of information flow across the frontal and visual brain areas (Goddard et al., 2016). We also used the same method in a passive object processing paradigm to investigate the spatiotemporal dynamics of information transaction between prei-frontal and peri-occipital areas and found that the peri-frontal and peri-occipital areas interacted, not only in the processing of categorical but category-orthogonal variations of objects (Karimi-Rouzbahani, 2018).

Here, in order to evaluate how (if at all) the task could influence the spatiotemporal dynamics of object processing in the human brain, we designed an object detection EEG study in which human subjects recognized target objects among a set of consecutively presented objects. Using, event-related potentials, frequency domain and multi-variate pattern analyses (MVPA) along with Granger causality, we made several observations: we saw that object categories were almost equally decodable from brain signals in both target and non-target conditions with late target-related differences across conditions on the whole-brain scale. Area-specific decoding across categories and across target/non-target conditions showed a significant initial target-related effect in prefrontal area. The target-related modulation was observed in turn in frontal, parietal, occipital and temporal areas suggesting a backward spread of task effect initiating in prefrontal areas, while the category information moved from anterior to posterior in the early window and from posterior to anterior brain areas in later windows. We also found the target-related information to be a better neural correlate for our behavioral object detection performance compared to the category-related information. These results provided deeper insights into the spatiotemporal dynamics of category- and task-related information processing in the brain.

## Methods

### Stimulus set

We used a subset of the well-known object image set of Kiani et al. (2007) which is freely available at (http://www.cns.nyu.edu/kianilab/Datasets.html#id4). The selected subset consisted of four categories of images namely ‘Animals’, ‘Faces’, ‘Fruits’ and ‘Objects’ each of which contained twelve exemplars (48 unique stimuli in total). The selection of images from the original image set was based on an exhaustive search to find exemplars with minimal differences in luminance, contrast and object area across the four mentioned categories. Some samples from the selected image set can be seen in Fig. 1A. Images had an area of 175 × 175 pixel and subtended around 8 degrees of visual angle when appeared on the screen during the experiment.

**Fig. 1.**
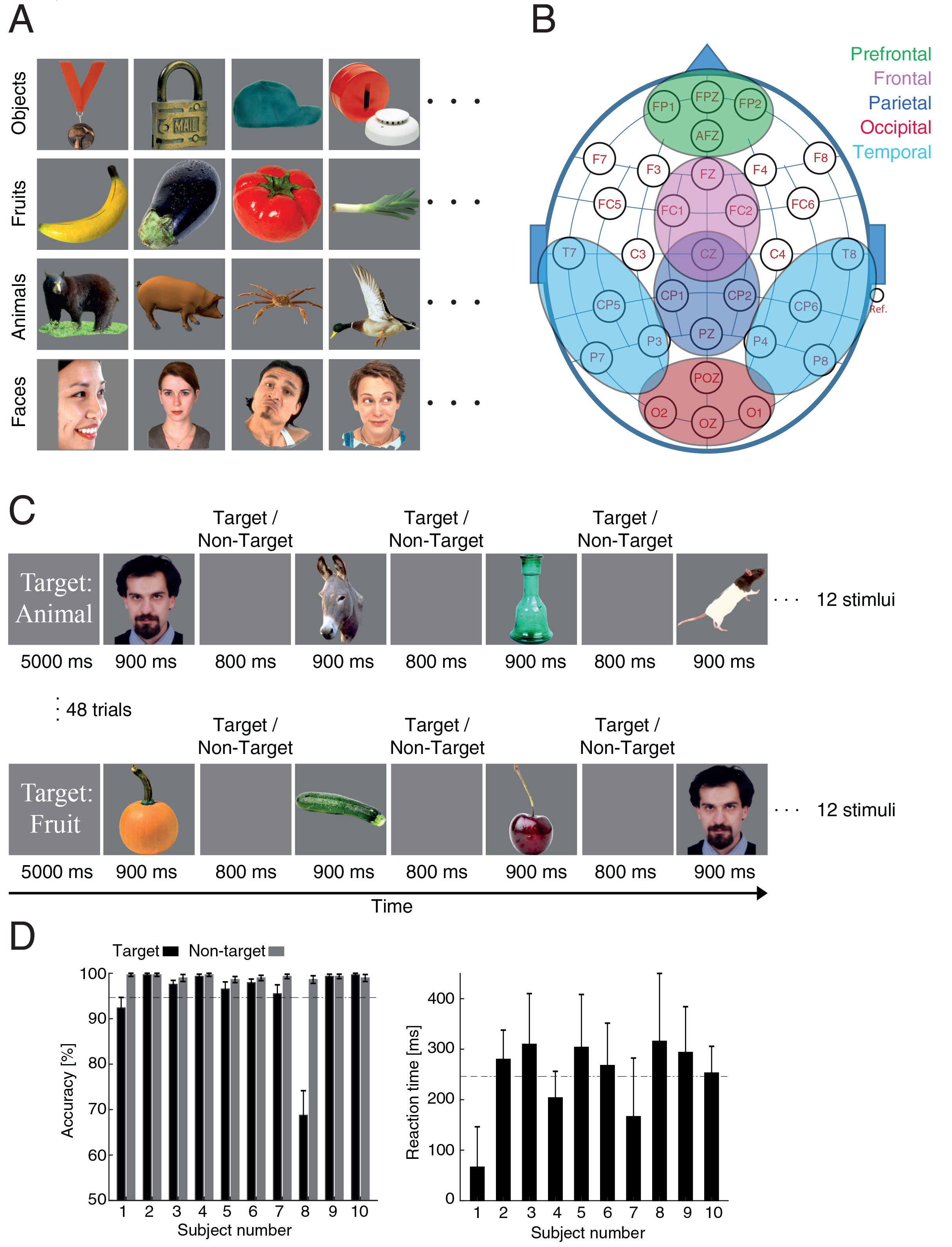
Image set, definition of areas, experiment description and behavioral results. (A), Four samples of the total twelve image exemplars within each of the four categories. The image set was obtained from (http://www.cns.nyu.edu/kianilab/Datasets.html#id4). (B), the definition of areas differentiated with colors. (C), Each trial of twelve stimuli started with the presentation of a text cue indicating the target category during the trial. The cue was presented for 5 s, followed by stimuli from target and non-target categories presented in random order for 900 ms with 800 ms of inter-stimulus interval. Subjects were asked to press the ‘space’ key after the offset of target stimuli. Each subject had 48 trials broken into 3 blocks with rest time in between the blocks. (D), Behavioral performance for each subject in detecting the target (black) and non-target (gray) stimuli. Error bars indicate the standard error across trials. The black dashed lines in the accuracy and reaction time plots show respectively the median accuracy and reaction times across all subjects’ data.

### Subjects

Twelve healthy volunteers participated in our single-session EEG recording experiment. The data from two subjects were removed from further analysis as a result of being noisy and artefactual. The remaining ten subjects (mean age of 25.4 years, eight right-handed, four females) had normal or corrected to normal vision. All subjects signed informed consents. The experimental protocol followed the guidelines of Helsinki’s declaration and was approved by the ethical review board of Shahid Rajaee Teacher Training University.

### Experimental design and task

Subjects participated in a single-session target-detection experiment which lasted for 40 minutes. They sat on a chair in a dimmed room facing a 35-cm away Asus VG24QE monitor on which the stimuli were presented. The presentation of the stimuli, as well as the recording of the responses, were done using Matlab Psychtoolbox (Brainard et al., 1997). Each trial began with the presentation of a cue for 5000 ms which indicated the target category to which the subjects had to respond (Fig. 1C). Specifically, subjects had to press the ‘space’ button of a computer keyboard (using their dominant hand) after the offset of the target stimulus and before the next stimulus appeared, which happened 800 ms later. We chose a go/no-go paradigm as compared to the two-alternative forced choice (2AFC) paradigm, as it was previously shown that the former could evoke a higher degree of target modulation (Bansal et al., 2014). Each trial included the presentation of twelve stimuli, half of which were the target category. The presentation of the target/non-target stimuli were in random order for each subject in each trial meaning that the six target stimuli could even appear consecutively. However, the frequency of the target compared to non-target categories was not significantly different in any of the twelve positions in the presentation order across subjects (Wilcoxon’s signed rank test, p < 0.05). Each unique stimulus was presented to each subject six times (three times as target) randomly across a total set of 48 trials. Accordingly, each subject was presented with a total of 576 (6 × 48) stimuli. The stimulus repetition was aimed at obtaining a higher signal to noise ratio in analysis. The stimuli were presented in three runs (each run included 16 trials), with 2 inter-run rest times each of which lasted for five minutes. The subjects were instructed to remain as stable as possible during the recording runs but were allowed to move their body during the rest times. They underwent a very short training session, on a different subset of stimuli, prior to the main experiment to become familiar with the task.

### EEG recordings and preprocessing

Electroencephalographic signals were recorded using a 32-channel amplifier with a sampling rate of 1000 Hz (eWave32; ScienceBeam Inc.; http://www.sciencebeam.com). The EEG amplifier also recorded the exact time instances of cue/stimulus presentation onsets using an optic sensor mounted on the corner of the monitor. The reference electrode was put on the right mastoid. We notch-filtered the recorded signals at 50 Hz to remove line noise and band-passed the signals in the range from 0.5 to 200 Hz respectively to remove signal drifts and high-frequency noise. A separate set of analysis were done with 0.05 to 100-Hz band-passed signals to ensure that no bias existed in the time course of the signals as the result of high-pass filtering (Widmann and Schröger, 2012; Tanner et al., 2015), which revealed no noticeable effect on the results. The filters were FIR filters with 12 dB roll-off per octave. We epoched (windowed) the signals around to the stimulus from −500 to +1500 ms relative to the stimulus onset. In order to remove artifacts from the signals, we decomposed the signals using Independent Component Analysis (ICA) as implemented in EEGLAB which used the runica algorithm (Delorme and Makeig, 2004). We selected the artefactual components using ADJUST plugin (Mognon et al., 2011). The ADJUST plugin, by extracting statistical spatiotemporal features from IC components, suggested the components which possibly reflected horizontal/vertical eye movements, eye-blinks and generic discontinuity in the signals. We also removed the artefactual stimulus-aligned signal epochs by visual inspection from further analysis. We did this artefact removal based on signal amplitudes: the epoch was removed from further analysis if it contained amplitudes greater than 4 × the standard deviation of the whole experiment. On average, 97.53 % of trials passed the artifact removal procedure and were used in the analyses (i.e. removed trials ranged from 7 to 24 across subjects). For the decoding analysis, the signals were smoothened using 5-ms moving average FIR filter and sampled every 5 ms.

### Multivariate decoding of categories and target effect

The decoding analyses were performed using Neural Decoding Toolbox (Meyers, 2013). Decoding of categories as well as the target effect (target vs. non-target conditions) were made across time using an LIBSVM classifier as in many previous studies (Chang and Lin, 2011). We also tested LDA and maximum correlation classifiers but did not observe any noticeable difference. Decoding was performed on each subject’s data separately across all possible pairs of categories and the two conditions. Accordingly, all six possible pairs of categories (animals vs. faces, animals vs. fruits, etc.) were decoded and their decoding results were finally averaged to obtain an average decoding curve for one subject. For the target effect, on the other hand, all the stimuli which were target were put against non-target stimuli and the decoding was performed across them. It should be noted that, the same set of stimuli (48 stimuli) was presented three times as target and non-target, in the experiment. Therefore, as the stimuli were the same across target and non-target conditions, this decoding reflected the target effect (i.e. the state of being the target vs. non-target). As mentioned above, the decoding was done at every 5-ms time instance across the epoch. The decoding procedure across categories for a single time instance was as follows: for a subject with no removed epochs, we had 144 epochs per category each of which had around 400 decoding time instance (every 5 ms from −500 to 1500 ms relative to stimulus onset). Ninety percent of epochs (90% × 144) from each of the two category were used to train the classifier and the remaining ten percent for testing the classifier. This was repeated for the other nine folds of the data and repeated 100 times for each fold to provide 900 classification accuracy values which were finally averaged to obtain a decoding value for a single time instance. This was repeated for and averaged across all pairs of categories to obtain a decoding value for an individual subject. This was repeated for every time point to obtain across-time decoding results. The same procedure was repeated for other subjects and averaged across them to obtain across-subject results. The decoding of target/non-target conditions was done in the same manner, but with target/non-target conditions replacing the category conditions.

### Granger causality analysis

In order to study the flow of information in the brain, we used a recently proposed version of Granger causality analysis (Goddard et al., 2016). As opposed to previous versions of Granger causality, which evaluated the flow of neural signals in the brain, the modified version, explicitly evaluates the flow of ‘information’ rather than unknown signals. In fact, in previous versions of Granger causality, the flow of neural signals was evaluated which might not necessarily have contained task-related information (e.g. category-or target-related information as in here). To that end, the desired information should be defined in the form of representational dis/similarities which are constructed for each area separately and compared for causal relationship across pairs of brain areas. In the following, we explain how the Granger causality analysis was implemented in this study.

Granger causality supports that time series ‘X’ can have a causal role in observing time series ‘Y’ if ‘X’ helps to predict the future values of ‘Y’ more accurately than when considering present values of time series ‘Y’ alone (Granger, 1969). Consider the case that frontal brain areas are considered to play a causal role in object category representations formed in occipital areas at a later time point. In that case, the representations of categorical information formed in frontal brain areas during the past time instances, if combined with the same-time past categorical information on occipital areas, can enhance the predictability of the present-time categorical information on occipital areas, compared to when predicting the present-time occipital categorical information based on its past values alone.

We did two analyses of Granger causality to study the spatiotemporal flow of both categorical information as well as target effects (across target vs. non-target conditions). For the categorical information, as was did previously (Goddard et al., 2016; Karimi-Rouzbahani, 2018), a representational dissimilarity matrix (RDM) was constructed, which contained the decoding rates (i.e. decoding was done as explained above with 6-fold cross-validation and 100 repetitions) across all possible pairs of stimuli. Accordingly, the dissimilarity matrix had a size of 48 × 48 (based on the number of unique stimuli) and was constructed separately for each of the five areas shown in Fig. 1B (i.e. the two temporal RDMs, which provided comparable results, were averaged and reported as ‘temporal’ in the following analyses). Next, we selected and reshaped the upper triangular elements of the RDM matrices (excluding the diagonal elements) into a vector (with 1104 elements, called ‘RDV’ here) and used it in Granger formulation as the across-category information vector.

For the target effect, on the other hand, we trained and tested the classifier with representations of exactly the same stimulus in target and non-target conditions. In other words, to obtain a single element of the desired target effect matrix, we classified the same stimulus in the target against non-target conditions. The result vector differentiated between being the target compared to non-target category (reflecting subject’s internal attentional set). This vector of target effect, which was later used in our Granger analysis, could represent the classical attentional modulation effects (e.g. such as multiplicative/additive modulation of tuning curves) generally investigated in neuronal studies (Moran and Desimone, 1985; Steinmetz and Moore, 2009). This vector, however, provided a high-dimensional modulation vector (in EEG space) as opposed to the classical one-dimensional attentional effect in neuronal studies. For more information about our Granger analysis method please refer to the two previous references (Goddard et al., 2016; Karimi-Rouzbahani, 2018). The category- and target-related representational vectors across each pair of areas were used in our Granger causality analysis separately using a partial correlation formulation as determined by (1):

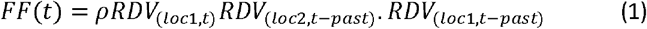

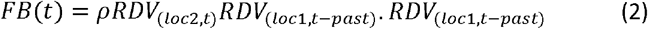

where *RDV*_(*loc*,*t*)_ is the representational dissimilarity vector obtained from the corresponding RDM on location lac at time *t* post-stimulus onset, and *RDV*_(*loc,t* −*past*)_ is the representational dissimilarity vector obtained by averaging the RDMs in a specific past window (e.g. from *t*-30 to *t*-60 ms relative to stimulus onset) on the same location. We chose the past time windows for the category- and target-related RDVs to be from −120 to −150 and from −30 to −60, respectively, for two reasons: first, the top-down object recognition and attentional feedback signals have respectively shown delays from around 10 ms (Gregoriou et al., 2009) to 140 ms (Lamme and Roelfsema, 2000) when transferring between anterior and posterior signals. Second, the effects (Figs. 5 and 5) were most significant in these chosen windows.

### Time-frequency analysis and cross-area coherence

In order to evaluate the time-frequency differences between the target and non-target conditions in different brain areas, we calculated the event-related spectral perturbation (ERSP) on every area by averaging the electrode signals within each area. This measure could provide an average time course index of relative changes in the EEG spectrum amplitude induced by the experimental events (Makeig, 1993) and was calculated using (3):

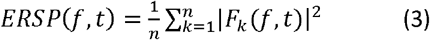

where *F*_*k*_(*f*, *t*) is the spectral estimate of trial *k* at frequency *f* at time *t* and *n* is the number of trials. This formulation, which was implemented in EEGLAB, used a time-varying amplitude spectra for each area using sinusoidal wavelets with increasing number of cycles (from 0.5 to 3 cycles) with frequencies and 50% overlapping data epochs. Here, the difference in ERSP maps are reported between the target and non-target conditions.

We also calculated cross-area coherence to help us understand whether (if at all) anterior and posterior brain areas operated in synchronized phases relative to the time of stimulus onset. The evaluation of cross-area coherence will also enable us to compare our results with previous studies which used cross-area coherence to evaluate the causal relationship between anterior and posterior brain areas (Summerfield et al., 2006). Cross-area coherence, which was calculated according to equation (4), provided a measure of phase-coupling which ranged between 0 (no phase-locking) to 1 (maximal phase-locking). The coherence of two signals, is essentially, their spectral cross-correlation normalized by signal powers. We used EEGLAB to calculate coherence across areas:

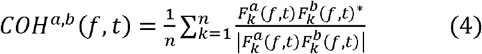

where * denotes the complex conjugation and *a* and *b* indicate the two areas; other notations were explained above. To obtain the signal from each area, we averaged the signals from electrodes within each area. Here, we reported the difference in COH maps between target and non-target conditions.

### Statistical testing

Statistical analyses were all performed in Matlab (version 2015b, The Mathworks, Natick, MA). In order to compare the subjects’ recognition accuracies and reaction times (Fig. 1D), we used the Wilcoxon’s signed rank test, and the significance level was considered to be p < 0.05.

To evaluate the significance of ERP signals at each time point (Fig. 2), we performed a Wilcoxon’s signed rank test between the target and non-target signals and obtained one p-value at each time point. Next, to correct the results for multiple comparisons, we applied an FDR correction (using Matlab mafdr function which used the Story’s method; Storey, 2002) and considered the final p-values as significant if they were lower than 0.05.

**Fig. 2.**
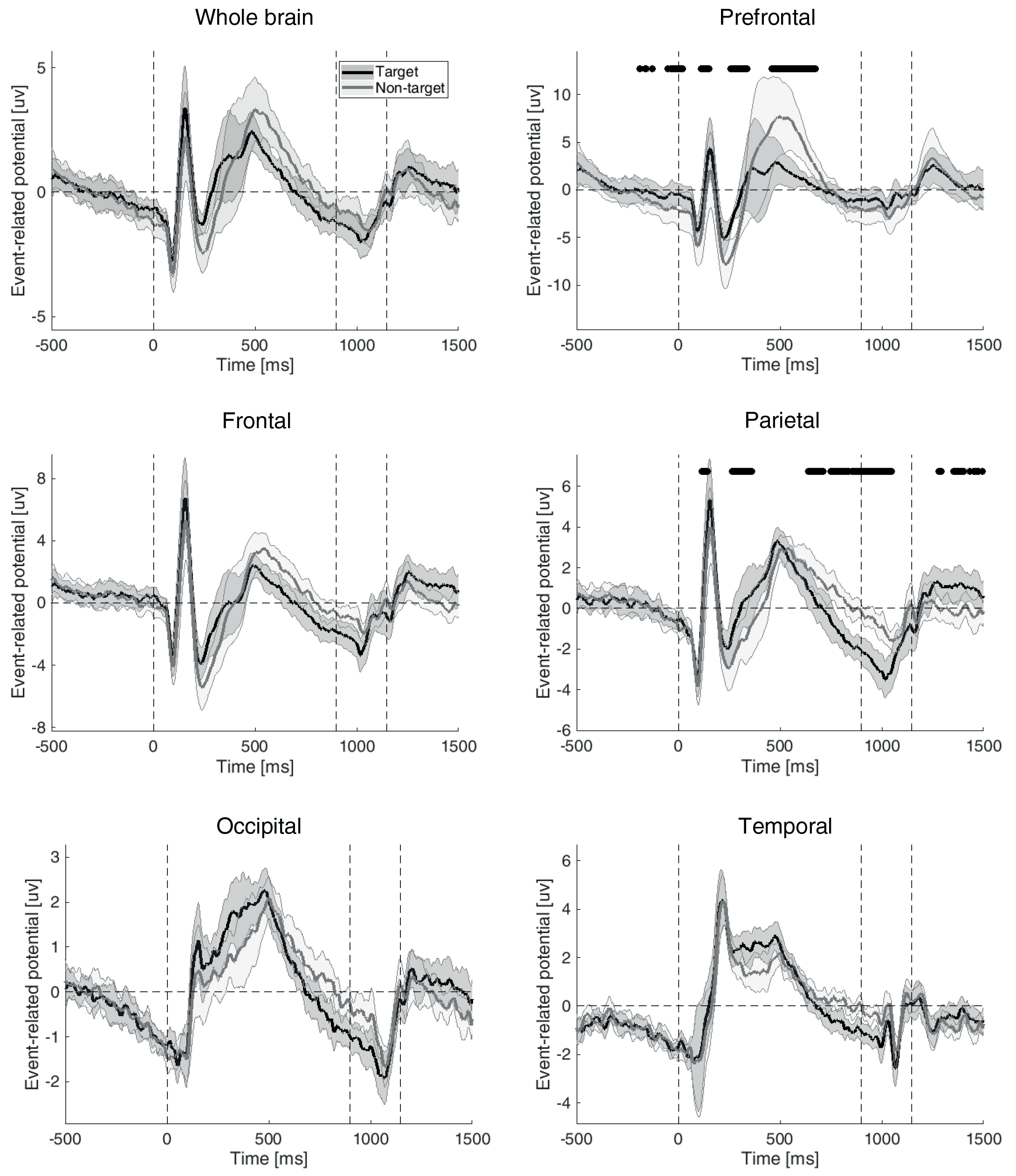
Event-related potentials recorded from the whole-brain and specific areas of the brain averaged across the subjects in target (black) and non-target (gray) conditions. The vertical dashed lines from left to right indicate the stimulus onset, offset and subjects’ median reaction times. The shaded error areas indicate the standard error across subjects and the black circles indicate the time points at which ERP values were significantly different between the target and non-target conditions (FDR-corrected Wilcoxon’s signed rank test).

To determine the significance of the time-resolved decoding curves (Figs. 3 and 4), we used a non-parametric bootstrap sampling with 1000 repetitions. More specifically, we shuffled the class labels (e.g. animals vs cars in categorical decoding and target vs non-target in target-modulation decoding) and decoded the signals 1000 times at every time point and obtained 1000 decoding values corresponding to the 1000 random subsample. Then, we compared the true decoding value (i.e. which was obtained based on true class labels) with the sampling-based decoding values and calculated the p-values based on one minus the proportion of sampling-based decoding values which were surpassed by the true decoding value. Finally, to obtain the time-resolved significant time points, we FDR-corrected the whole set of (601 time points) p-values and considered the true decoding values as significant whenever their corresponding p-values were smaller than 0.05.

**Fig. 3.**
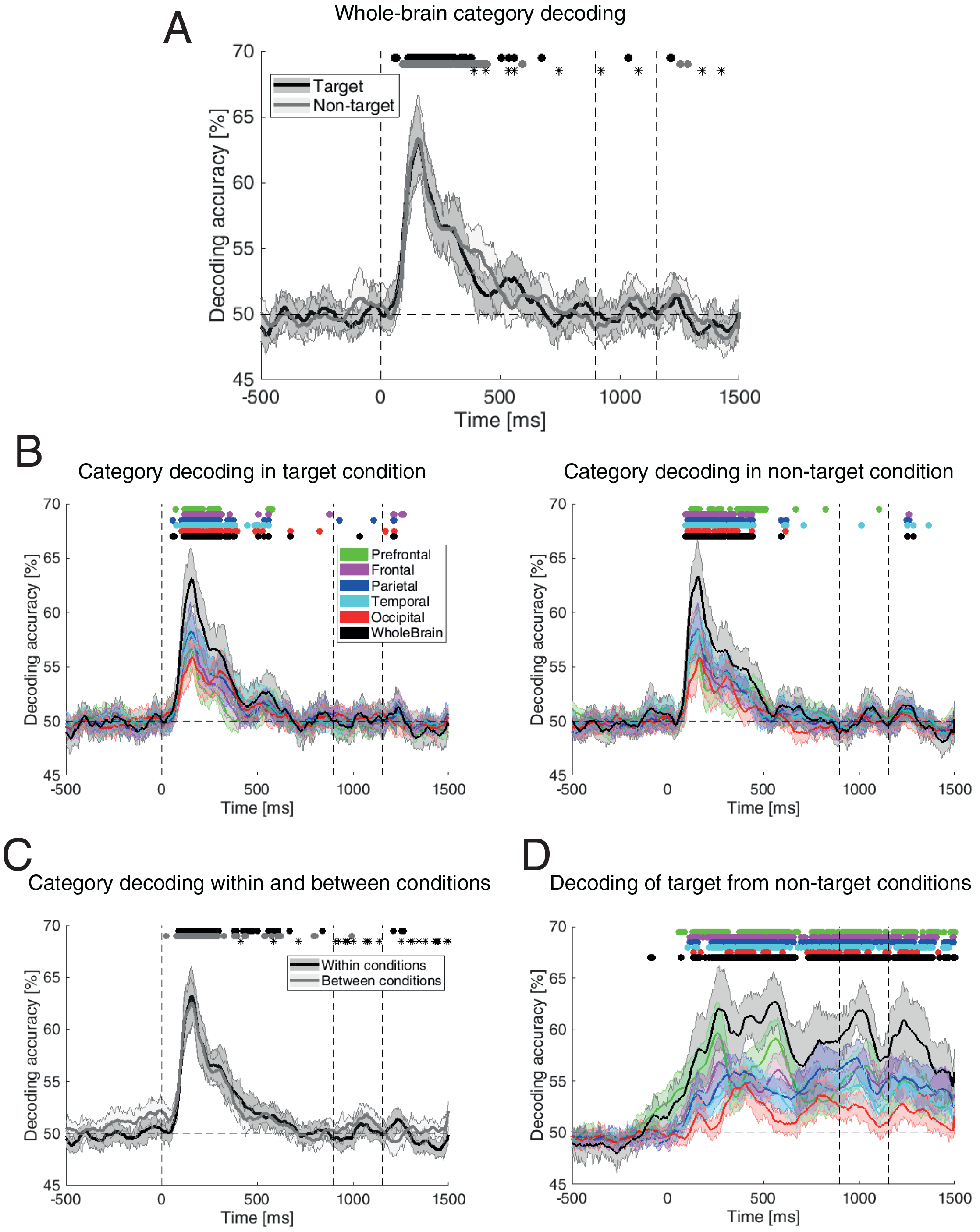
Multivariate pattern decoding of category and target-related information in the brain. (A), Decoding of categories in the target and non-target conditions using all 31 electrodes on the head. (B), Decoding of categories in the target (left)/non-target (right) condition on the whole brain and in each area. (C), Decoding of categories within and between target and non-target conditions. (D), Decoding of conditions (target vs. non-target) on the whole brain as well as on each area. The circles and asterisks indicate respectively the time points at which the corresponding decoding curve was significantly different from chance and the time points at which the decoding curves were significantly different across target and non-target conditions (FDR-corrected Wilcoxon’s signed rank test). The shaded error areas indicate the standard error across subjects. The horizontal dashed line indicate chance decoding of 50% and the vertical dashed lines from left to right indicate the stimulus onset, offset and subjects’ median reaction times.

**Fig. 4.**
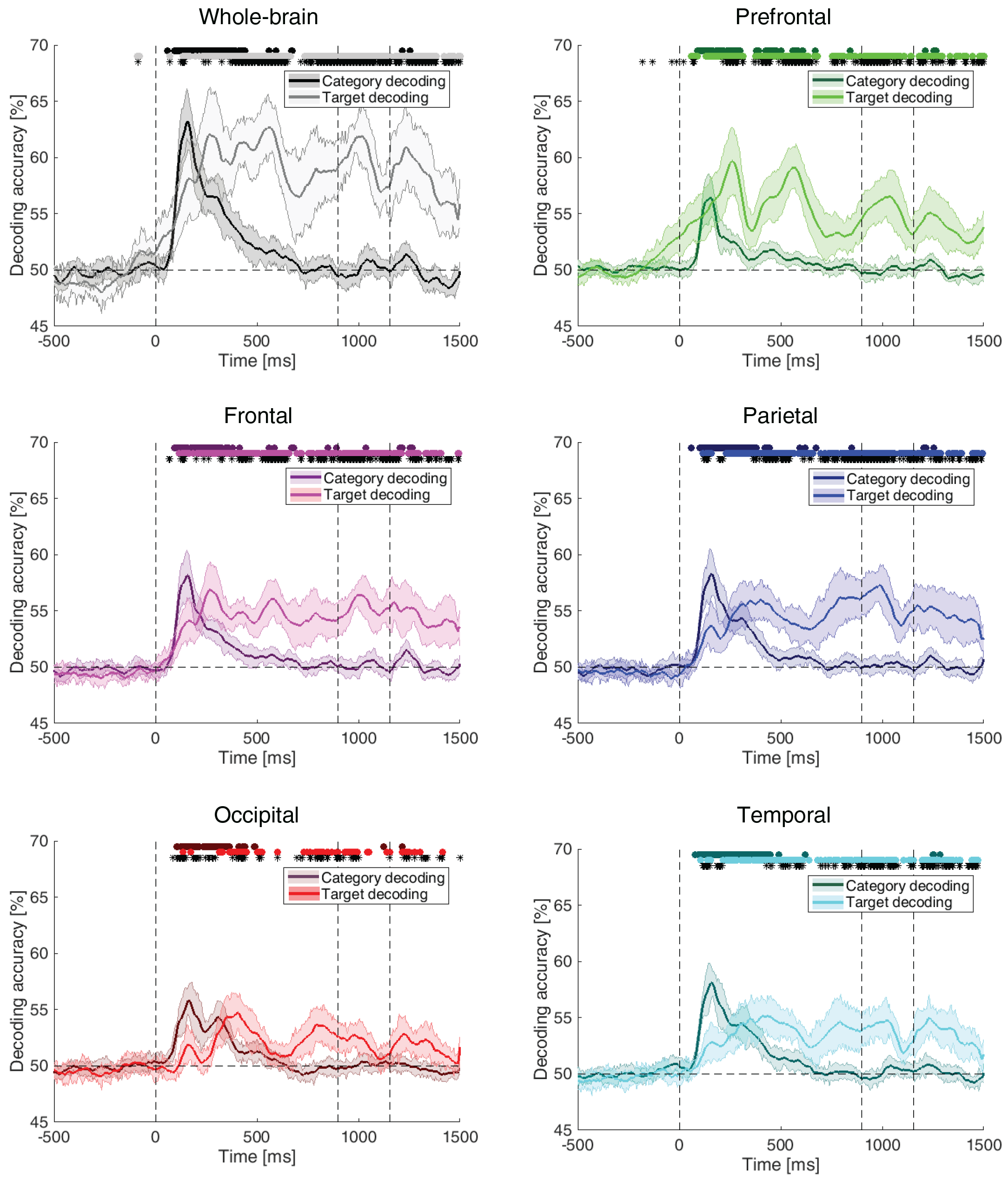
Comparison of category- and target-related information decoding. These results are extracted from the results shown in Fig. 3. Accordingly, to obtain the category decoding results, the target and non-target results of each area (Fig. 3B) were averaged and plotted over the target-related decoding results (Fig. 3D). Asterisks indicate the time points at which the category- and target-related decoding curves were significantly different (FDR-corrected Wilcoxon’s signed rank test). Other details are the same as in Fig. 3.

The significance of partial correlations (movement of information on the head) was also evaluated using bootstrap sampling with 1000 repetitions (Figs. 5 and 7). Specifically, 1000 random partial correlations were calculated at each time instance by shuffling the RDM elements of both areas. Then the difference in partial correlations was calculated between the feedforward and feedback (feedforward-feedback) flows of information (respectively obtained from equations (1) and (2)) from the true as well as from the randomly shuffled RDMs. Then, the difference in partial correlations obtained from the true RDM was compared with those obtained after shuffling the RDMs: the p-values were calculated as one minus the proportion of differences with lower values than the true partial correlations difference. Finally, the p-values were corrected for multiple comparisons (using Matlab mafdr function) and considered significant if they were smaller than 0.05 (p<0.05).

**Fig. 5.**
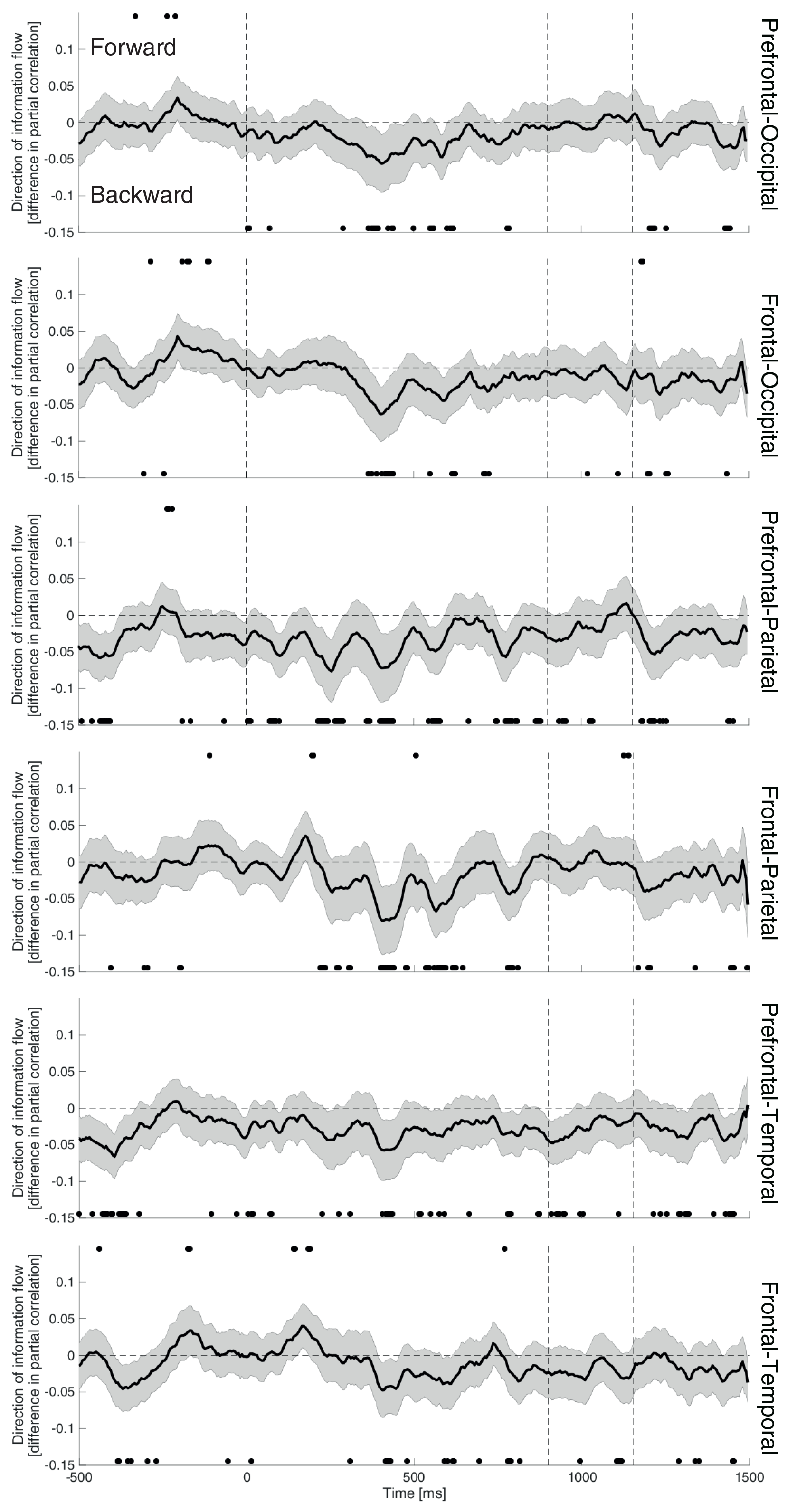
Flow of target-related information on the head across different areas. Results of difference in (feedforward minus feedback) partial correlations (as explained by equations (1) and (2)) are provided across different areas. Positive and negative values indicate respectively the feedforward and feedback of information flow across the indicated areas. Significance of the results was evaluated using random bootstrapping (FDR-corrected p-values with p < 0.05 as significant) as explained in the text and significant time points are indicated with black circles.

To evaluate the significance of ERSP and cross-area coherence maps, we used the same bootstrap sampling of target vs non-target trials with 1000 repetitions. Specifically, we obtained 1000 random ERSP and cross-area coherence maps by randomly assigning the trials to the target and non-target conditions. We compared the true ERSP and cross-area coherence maps (Supplementary Figs. S1B and S3) with those obtained from random bootstrapping and considered true value at specific time-frequency point as significant if it provided p < 0.05 after multiple comparison correction applied on the whole matrix.

The p-values of correlations between the decoding and behavioral results (Fig. 8) were obtained from Pearson linear correlations and were corrected for multiple comparisons before deciding on their significance (which had a level of p < 0.05).

## Results

This study was designed to evaluate the spatiotemporal dynamics of category and task-related information processing in the brain during the recognition/detection of target (attended) objects in a sequence of object images. To that end, ten human subjects participated in an object detection experiment in which their EEG signals were recorded. Subjects were quite fast (median reaction time = 246 s) and accurate (average recognition accuracy = 94.65%, SD = 9.38%) at performing the task, which shows that they were well-acquainted with the task and attentive during the experiment (Fig. 1D). Because of the high behavioral performance of subjects in this experiment, the analyses were limited to correct trials (the results of all trials also showed similar results).

### Comparison of (ERP/ERSP) signals in target versus non-target conditions

In order to compare possible differences between the stimuli’s evoked potential in target and non-target conditions, we calculated event-related potentials in those conditions. As the stimuli were the same in both conditions, differences in their ERPs could be associated with subject’s internal information (e.g. categorical attention). We plotted the ERPs from −500 to +1500 ms aligned to the stimulus onset on the whole brain as well as the each of the five areas separately (Fig. 2). In order to be able to interpret the ERP signals we need to have prior information about different time stages of signal processing. In the following paragraph, we will explain what brain processes are different time-windows of the analyses associated with.

The window from −500 to 0 ms, could dominantly contain preparatory information about the target category (categorical attention) rather than response-related information from the preceding trial. This is justified by the fact that only less than 25% and 8% of the reaction times were registered in the window from the stimulus offset to +300 ms and +400 ms post-stimulus offset (which is from −800 to respectively −500 and −400 ms relative to the onset of the current stimulus; Supplementary Fig. S1A). Moreover, the window from the stimulus onset to its offset (0 to +900 ms), can be split into three main overlapping phases according to previous studies: the first phase, which is mainly associated with visual processing starting from the stimulus onset to around +400 ms post stimulus onset (Hebart et al., 2018; Isik et al., 2013; Kaneshiro et al., 2015), followed by a second phase, associated with decision-making (VanRullen and Thorpe, 2001; Kelly and O’Connell, 2013), which is probably ongoing within the window from +150 ms to +800 ms and a third phase, associated with motor command preparation (Thorpe et al., 1996), which covers the window from around +200 ms and lasted to the end of stimulus presentation time at +900 ms or after the stimulus offset. The motor action (button press) was recorded after the stimulus offset, based on which the behavioral outcomes were measured. Please note that, as opposed to what is normally done in ERP analyses (Thorpe et al., 1996), we did not remove the pre-stimulus baseline activities from the ERPs as this could remove possible pre-stimulus target-related effects which could appear as baseline biases.

While the ERPs showed observable differences between the target and non-target conditions on the whole-brain analysis plot (Fig. 2, top-left panel) especially after 100 ms, the FDR-corrected significance testing (Wilcoxon’s signed rank test) showed no significantly different ERP values across the time points between the target and non-target conditions. The ERPs of the target condition, seem to show an earlier upward and downward trends compared to the non-target ERPs especially in the window from 200 to 1000 ms which might reflect an earlier processing initiation for the target compared to the non-target conditions. Looking for target-related effects across different brain areas (Fig. 2), we saw that significant target-related effects (p < 0.05; FDR-corrected Wilcoxon’s signed rank test) were only observable on prefrontal and parietal brain areas. Interestingly, while the parietal areas showed the effect during the post-stimulus windows in four dominant time spans (namely from 115 to 143, 267 to 359, 638 to 1047 and from 1282 to 1500 ms), the prefrontal areas showed a much earlier effect even before the onset of the stimuli (starting at around −190 ms). As supported by the literature, the target-related effects, which were observed on the parietal cortices, could possibly be mainly attributed to decision-related processes (Kelly and O’Connell, 2013). These processes are proposed to be mainly undergone by the multiple demand (MD) network spanning from the frontal to parietal cortices (Woolgar et al., 2015a; Jackson et al., 2017). On the other hand, while the earlier windows (e.g. from 0 to 100 ms) of post-stimulus effects observed on prefrontal areas can reflect the involvement of orbitofrontal cortex in the processing of low-frequency spatial contents of target (Bar et al., 2006; Thorpe et al., 1983; Foxe and Simpson, 2001; Karimi-Rouzbahani, 2018), the later windows (e.g. from +150 ms) can also be attributed to decision- and motor-related processes (Duncan and Owen, 2000; Daliri et al., 2013). The pre-stimulus effects can be associated with predictive signals to prepare the visual areas to detect the target/ignore the non-target category of the coming stimulus (Summerfield et al., 2006; Battistoni et al., 2017; Stokes et al., 2009; Soon et al., 2013). This is explained in more details in Discussion. The earlier temporal dynamics of target ERPs compared to non-targets can also be observed on almost all areas (Fig. 2).

We also calculated the event-related spectral perturbations (see Methods) to compare the event-locked time-frequency dynamics across the target and non-target conditions (Supplementary Fig. S1B). In order to be able to compare the two conditions more explicitly, we provided the difference time-frequency matrices which were calculated as the ERSP in target minus the ERSP in non-target condition. The whole-brain difference ERSP matrix (top-left panel) showed significant reduction of power mainly in the delta (0.5-4 Hz), theta (4-8 Hz) and alpha (8-13 Hz) bands starting at 180 ms post stimulus which could be associated with decision making, attention and motor processes (Spyropoulos et al., 2018; Nacher et al., 2013; Rohenkohl and Nobre, 2011; Davis et al., 2012), respectively. While the whole-brain result was mainly reflecting the prefrontal reduction of power starting at around the same time instance, other regions showed area-specific effects as follows. The occipital brain area showed a reduction in alpha and theta bands power which are generally associated with attentional processes affecting this area after 450 ms (Klimesch, 2012; Rohenkohl and Nobre, 2011). As the early visual processing was reflected in both target and non-target ERSP maps (data not shown), the difference ERSP map, which represented their differences, reflected only the effects during post-sensory processing times. The temporal areas, also showed a delta and theta power reduction associated most probably with motor preparation (Davis et al., 2012) which might have leaked from parietal and frontal brain areas. The frontal and parietal brain areas, showed an early (between 500 to 920 ms) as well as late (after 1300 ms) ~20 Hz power reduction effects previously associated with motor actions (Davis et al., 2012). Altogether, these results suggested that the power of the signals in different brain areas provided timely-ordered evidence for attention-, decision- and motor-related signals which were expected to be different across the target and non-target conditions based on our paradigm design. Specifically, earlier time windows (e.g. covered by the window from 200 to 1200 ms) showed significant target-specific effects in many areas, followed by motor signals in parietal and frontal brain areas.

### Decoding of category- and target-related effects

The ERP and time-frequency results which were shown above, although valuable, remained silent on the spatiotemporal dynamics of categorical and target-related (attentional) information in the brain. More specifically, we could not decide what information the observed ERPs were carrying; it was unknown whether these brain signals contained information about categories or the task (stimulus being target/non-target). Therefore, we used multi-variate pattern decoding to investigate the information content of the EEG signals as explained in Methods.

Although the whole-brain time-resolved categorical decoding results for the target and non-target conditions showed significant differences at several time instances (see the asterisks in Fig. 3A), especially after 300 ms which is on par with previous results on the task effect at around the same time windows (Hebart et al., 2018), they showed largely similar patterns throughout the analysis window (Fig. 3A). Importantly, while the target condition showed a decoding curve which reached significance at an earlier time point (60 ms compared to 95 ms obtained for the non-target conditions), the non-target condition showed continuously significant decoding values which lasted for more extended windows (until 445 ms) compared to the target decoding condition (until 380 ms). The temporal dynamics of the decoding curves are in agreement with a rich literature on the temporal dynamics of object processing in the brain (Karimi-Rozubahani et al., 2017a; Kaneshiro et al., 2015). The decoding results are provided for each area separately in Supplementary Fig. S2A. While the decoding curves on prefrontal areas showed windows of significant difference between target and non-target conditions (e.g. from 370 ms to 575 ms and from 700 to 785 ms), other regions showed this effect only at some sparse time points.

Next, in order to investigate which brain areas contributed more to the processing of categorical information, we decoded the categorical information using area-specific electrodes (Fig. 1B) separately for the target and non-target conditions (Fig. 3B). The decoding curves repeated many of the whole-brain results: the target decoding curves at prefrontal, parietal and temporal areas reached significance at earlier times compared to the non-target decoding curves (respectively with 50, 55 and 15 ms of delay in non-target compared to target condition, with an opposite effect for the frontal area with 25 of delay in target condition and no change in timing at occipital areas). Therefore, it seems that the state of being the target category during object recognition, can speed up the processing of objects, especially at areas previously associated with top-down attention (Woolgar et al., 2015a).

The decoding results provided above, which showed many aspects of similarity between the target and non-target conditions, suggested that these conditions might have provided similar object representations despite their relevance to subjects’ goal (i.e. being the target of an object recognition task), as was previously observed (Hebart et al., 2018). In order to test this hypothesis, and to see whether the object representations of one condition (e.g. target) could be generalized to the opposite condition (e.g. non-target) we performed a cross-condition decoding. To that end, we trained the decoding classifier with representations in one condition and tested it with those of the opposite condition; so there were two decoding rounds, one with the target and the other with the non-target condition as the training set and the other as the testing set. The results of the two decoding rounds were finally averaged to obtain these results which we called the “between-condition” decoding results and compared them to those obtained from the “within-condition” results which were obtained by averaging the decoding results from target and non-target conditions reported in Fig. 3A. Results showed that, as previously observed across a set of different tasks (Hebart et al., 2018), the state of being the target or non-target in a detection/recognition task had an ignorable effect on the representations of objects decoded from scalp activities (Fig. 3C). This was true for every individual area of the brain (Supplementary Fig. S2B). This was unexpected, however, as we observed a significant amount of target-related information in the ERPs (Fig. 2) and ERSPs (Supplementary Fig. S1B) explained above. Therefore, we suspected that, our decoding analyses might have been unable to capture the task information from the object representations.

In order to rule out the above possibility, and to see whether the target-related information could be accessed by the same decoding procedure as was used for categorical information, we decoded the target stimuli from non-target stimuli without considering their assigned categories (Fig. 3D). Interestingly, results showed significant decoding values for the whole-brain signals throughout the decoding window. The decoding curve experienced two bumps of information at around 260 and 550 ms and remained significant until the end of analysis time. According to the experimental setup, and the ERP effects explained above (Fig. 2), the whole-brain decoding curve could be associated with an early target-related, an intermediate-time decision-related and a late motor-execution components with descending amplitudes. Next, we investigated the target-related effect on separate brain areas (Fig. 3D, different colors). The highest-to-lowest decoding values (averaged during the −200 to 900 ms window) were observed at the prefrontal, frontal, parietal, temporal and occipital areas (the average decoding values were respectively 54.7%, 53.32%, 53.26%, 52.77% and 51.63%). The higher involvement of prefrontal, frontal and parietal brain areas compared to the occipital and temporal areas in the processing of subject task is on par with previous results suggesting the involvement of the multi-demand (MD) network in the processing of tasks and top-down attention (Jackson et al., 2017; Woolgar et al., 2015b). Very interestingly, the prefrontal areas showed an above-chance decoding value which started to rise from chance prior to the stimulus onset and reached significance (p < 0.05, FDR-corrected bootstrap sampling test) at around 60 ms post-stimulus, earlier than any other area. This pre-to-post stimulus effect, which was also reflected on the whole-brain result, was not observed on any other areas. The target-related effects then appeared on temporal, frontal, parietal and occipital areas respectively at 105, 115, 115 and 135 ms post-stimulus onset. This early rise of target-related information on prefrontal areas seems to work as a prediction signal generated on prefrontal cortices, as previously suggested to be constructed in prefrontal cortex and sent back to the fusiform face area (FFA) for face recognition (Summerfield et al., 2006). However, more evidence is needed to support this proposal, which is provided below.

### Comparing the spatiotemporal dynamics of categorical and target-related information

The target-related information processing which showed distinguishable rates and dynamics across brain areas (Fig. 3D) suggested distinct roles for different brain areas in the processing of subjective top-down versus sensory-driven bottom-up mechanisms of the brain in object recognition, which needed further assessment. For that purpose, we compared the patterns and the dynamics of categorical and target-related information on the whole-brain scale as well as on individual areas (Fig. 4). Please note that, for the sake of comparison, and as in above, the reported categorical decoding values are averaged across (all six) pairs of categories; therefore, the chance level is at 50% (Fig. 4). The category decoding results of Fig. 4 are obtained by averaging the target and non-target decoding curves of Fig. 3. The whole-brain results showed comparable decoding rates for category- and target-related information with them reaching significance at +60 ms and −90 ms relative to the stimulus onset time. The level of category-related information significantly surpassed the target-related information at −85 ms (see the asterisks). The reason for the pre-stimulus target-related information observed in prefrontal areas is because some subjects could predict encountering target/non-target stimuli in the coming trials, as a result of unbalanced distribution of target/non-target conditions across their trials’ stimuli. This is explained further in Discussion. As mentioned above, the precedence of target-related information compared to category-related information seem to be initiated in prefrontal areas (Fig. 4, top-right panel) and serve as a top-down excitatory/inhibitory signal (Summerfield et al., 2006). In fact, the prefrontal area was the only region which revealed pre-stimulus task-related information, with a significantly higher decoding values for the target-related (task-evoked) compared to category-related (category-evoked) information in most pre- and post-stimulus time instances. While the category-related decoding curves peaked at around 160 ms, the target-related information peaked at around 265 ms post-stimulus.

The absolute amplitude of target-related decoding curves and their amplitudes relative to the category-related decoding curves decreased by going from prefrontal to frontal, parietal, temporal and occipital areas. While the target-related decoding curve on prefrontal and frontal areas experienced a descending trend after their first peaks which occurred before the stimulus offset, the parietal, temporal and occipital areas showed an ascending pattern. These results suggested that there might be a backward movement of target-related information in the brain initiated in prefrontal/frontal cortices which reached category-processing cortices such as parietal, occipital and temporal areas to modulate the processing of target category.

### Flow of target-related information in the brain

To quantitatively test the possible movement of information from anterior to posterior brain areas, we performed Granger causality analysis to evaluate the flow of target information in the brain (see Methods for the details of this analysis). Results showed that both the prefrontal and frontal brain areas sent target-related information to the posterior regions of the head (Fig. 5) mainly from 0 to +400 ms (which peaked at around +400 to +500 in almost all areas) whose amplitude declined in the window from +500 to +900 ms relative to the stimulus onset. Specifically, while the information movement showed some sparsely distributed time points of significant backward/forward flow in the pre-stimulus time windows, the movement curves showed many continuously distributed significant time windows indicating the flow of target-related information towards the back of the head in the window from 200 to 800 ms relative to the stimulus onset. As these flow analyses were obtained from a delay window of 40 ms across the anterior and posterior brain areas, we could conclude that the attentional effects may have passed between two to eight synapses (Azouz and Gray, 1999) to reach the visual processing areas of the brain which seems to be longer than previously reported results (8-13 ms) for top-down attentional flow for non-human primates (Gregoriou et al., 2009) and on par with the speed of visual information flow across anterior and posterior brain areas in humans (Goddard et al., 2016). These results supported the flow of target-related signals from anterior to posterior brain areas in object recognition.

In order to obtain a deeper insight into the spatiotemporal dynamics of target-related information flow in the brain, we conducted a univariate searchlight decoding between target and non-target conditions on every individual electrodes and interpolated their results for scalp-decoding topographies to trace the target-related information on the head (Fig. 6). Specifically, as we previously did (Karimi-Rouzbahani, 2017a), compared to the decoding curves in Fig. 3D, which were obtained using all the electrodes simultaneously (31-dimensional space), these results were obtained using an individual electrode at a time (1-dimensional space) before finally interpolating them for whole-brain scalp topographies. Please note that the decoding topographies are obtained by averaging the time-resolved decoding values in 50-ms windows around the indicated time points. As the results show (Fig. 6), we see above-chance decoding on prefrontal areas in 0 to 100 ms windows, which extended from prefrontal to frontal, central and parietal areas in the windows from 150 to 350 ms. These results seem to reflect the dynamics of backward target-related (attentional/predictive) information flow previously explained for Fig. 5. On 350 ms and the coming windows up to 1000 ms, the decoding seem to have covered the parietal and central brain areas as well as the frontal regions of the brain which might reflect decision-related information being processed in the frontoparietal (MD) network (Woolgar et al., 2015b). This decision-related process can also be supported by the change in the delta-band (0.5 to 4 Hz) coherence across the posterior and anterior brain areas (Nacher et al., 2013) which was also observed here in our cross-area coherence analysis (Supplementary Fig. S3). The following windows (from 1000 ms onward), which showed target-related information mainly concentrated on centrofrontal areas, seem to be representing the response-related activity in motor cortex. Altogether, these topographical target-related decoding maps provided a systematic spatiotemporal flow of information in the brain, which supported previous results on the functional role of sensory- and task-related brain areas in object recognition and action.

**Fig. 6.**
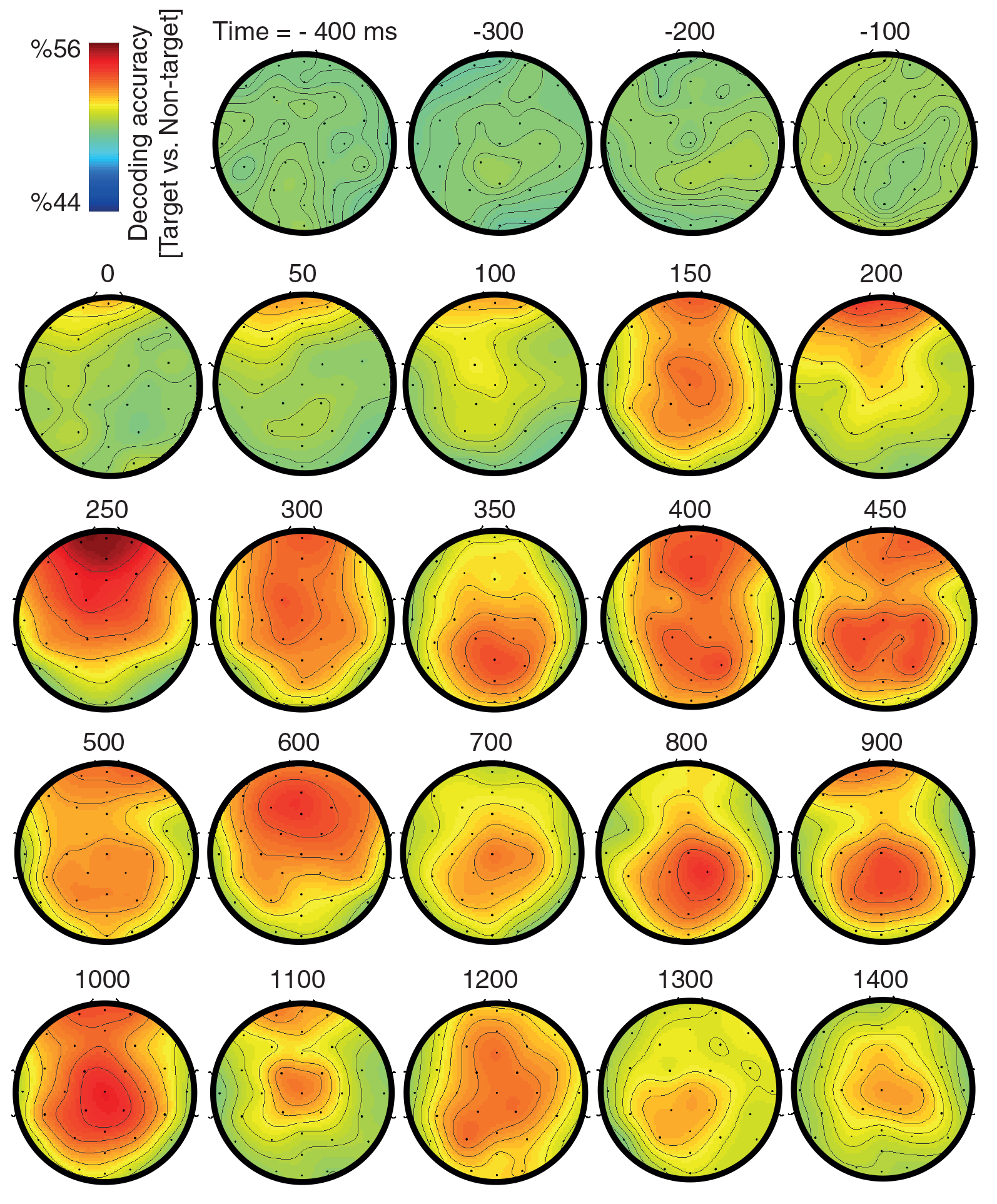
Topographic decoding maps showing target-related information movement on the head. These maps were generated by decoding the conditions (target vs. non-target) on each electrode separately and finally plotting their superposition on the scalp. The decoding values for between-electrode areas were calculated by interpolations implemented in EEGLAB. The reported decoding values are averaged in the span from −25 m to +25 ms relative to the indicated time points.

### Flow of category-related information in the brain

Using a similar Granger causality analysis, two previous studies (Goddard et al., 2016; Karimi-Rozubahani, 2018) have observed the flow of categorical information from the peri-frontal areas of the brain towards the peri-occipital areas during object processing. As our previous study (Karimi-Rozubahani, 2018), which used a passive object viewing paradigm (probably resembling current study’s non-target condition), showed a different timing of categorical information flow compared to the seminal method-developing study (Goddard et al., 2016), with an active object recognition paradigm (probably resembling current study’s target condition), it was interesting to know whether the category-related flow of information differed in the target versus non-target conditions of the current study. Therefore, we evaluated the flow of category-related information in the brain across anterior and posterior brain regions (Fig. 7). While the non-target condition showed significant directed information flow only at some sparsely distributed time points, significantly backward and forward information flows were observed respectively at around the 230 to 260 ms and 440 to 460 ms windows across the prefrontal- and frontal-occipital pairs in the target condition (Fig. 7). Other anterior-posterior pairs did not show continuously positioned significant forward/backward flows. These results showed that the category-related information had a different temporal patterns compared to the target-related information dynamics explained in Fig. 5: while the target-related information, which probably represented the top-down attentional modulation (Paneri and Gregoriou, 2017), showed an anterior-posterior movement of information, the category-related information, which explicitly reflected categorical information, showed an earlier anterior-posterior and a later posterior-anterior direction of categorical information movement in the brain. Please note that the delay which was used in the Granger causality analysis (equations (1) and (2)) for the target- and category-related information were respectively 40 and 150 ms. This suggested an earlier (starting at around 80 ms) movement of category-related information from anterior to posterior brain areas probably from orbitofrontal to occipital cortices for category representational enhancement (Bar et al., 2006; Karimi-Rouzbahani, 2017c), and a later (starting at around 300 ms) posterior to anterior movement probably from lateral occipital to frontal cortices for decision-making purposes (DiCarlo et al., 2012; Karimi-Rouzbahani, 2017c).

**Fig. 7.**
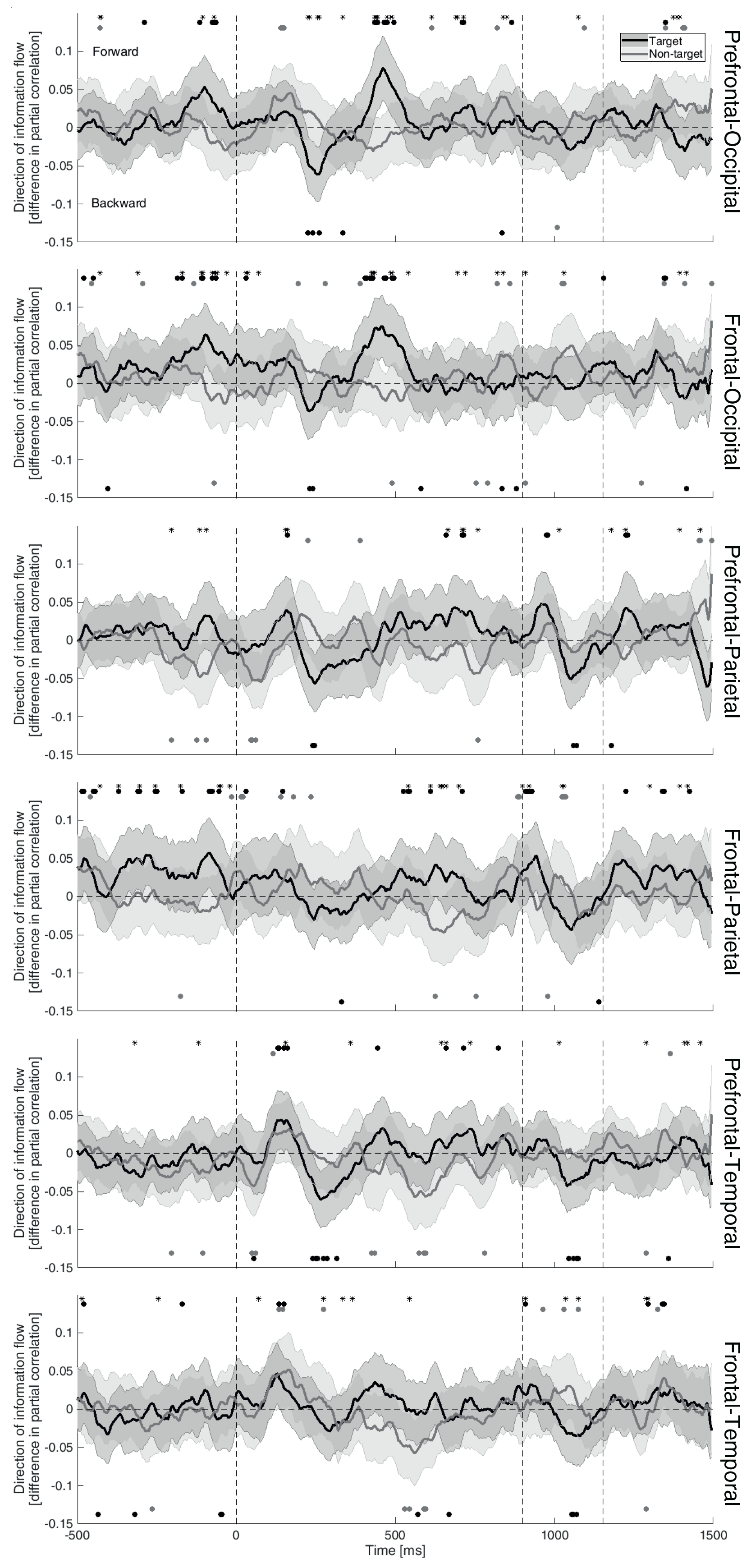
Flow of category-related information on the head across different areas for the target (dark) and non-target (gray) conditions. Results of difference in (feedforward minus feedback) partial correlations (as explained by equations (1) and (2)) are provided across different areas. Positive and negative values indicate respectively the feedforward and feedback of information flow across the indicated areas. Significance of the results was evaluated using random bootstrapping (FDR-corrected p-values with p < 0.05 as significant) as explained in the text and significant time points are indicated with corresponding colors. Asterisks show the time instances at which the target and non-target conditions showed significantly different values (FDR-corrected Wilcoxon’s signed rank test).

### How much can the brain representations explain behavioral results?

After making several suggestions in previous sections about the associations between different stages of decoding and their corresponding behavioral output, we did a correlational analysis to provide quantitative supports for our claims. For that purpose, we calculated the Pearson linear correlation between the category and condition (target vs non-target) decoding vectors across time (Fig. 4) and the behavioral vectors (i.e. either object recognition accuracy or reaction times (Fig. 1D)). The mentioned ten-element vectors consisted of decoding/behavioral results each of which obtained from one individual. We have previously suggested that the final windows of decoding (probably after 400 ms) reflected decision-making and response-related signatures. Accordingly, the whole-brain results showed a rising trend of positive correlation between the behavioral accuracy and the condition decoding after 500 ms which showed significance after the stimulus offset, and a falling-to-negative correlation trend between the subjects’ reaction times and the condition decoding which also showed several significant time points before and after the stimulus offset (Fig. 8A, top-left panel). The whole-brain results seem to have been mainly driven by the frontoparietal network, which have been previously associated with decision-making and motor actions (Fig. 8A, second row from top), rather than occipital and temporal brain areas. These results are interesting in that they showed positive and negative correlations between the brain decoding of conditions and respectively the subjects’ recognition accuracy and reaction times, supporting the behavioral relevance of our condition decoding analysis. However, these correlations were almost absent between the category decoding and the behavioral accuracy and reaction times (Fig. 8B). Specifically, while category decoding and recognition accuracy/reaction time showed several points of negative/positive (from 0 to 550 ms) and positive/negative (from 550 to 900 ms) correlations, it showed a change of trend after the stimulus offset, which is the most expected time window for brain-behavior correlation. Therefore, it seems that the condition decoding seem to provide a more accurate neural correlate for our object recognition/detection task. Specifically, rather than performing categorization, subjects seem to have concentrated on the detection of target from non-target stimuli.

**Fig. 8.**
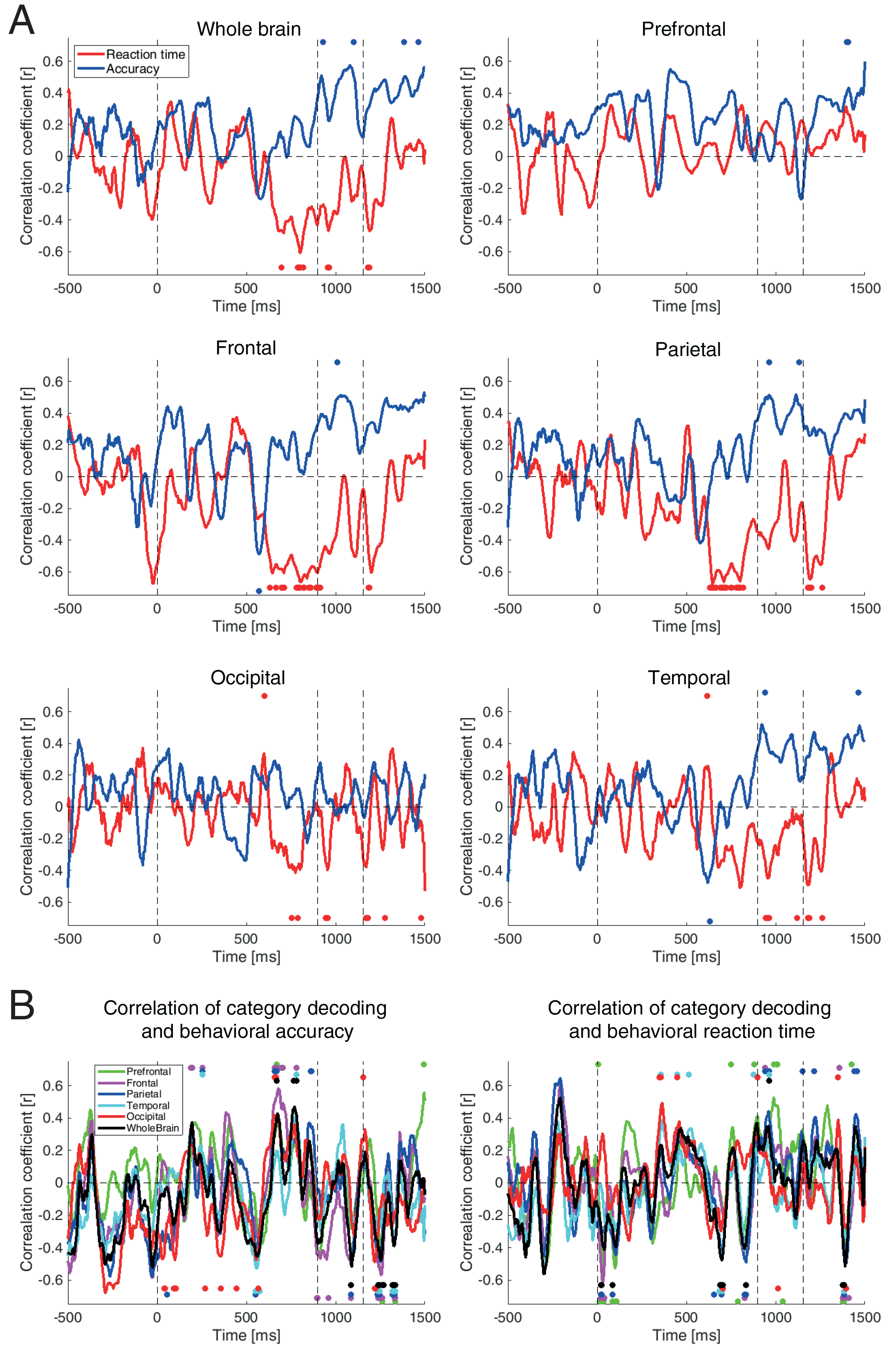
Correlation between target- and category-related results and behavioral performance. To obtain these graphs, we calculated the correlation between the behavioral results and the decoding vectors (as obtained from the ten subjects’ data). (A), Red and blue lines indicate the correlation between the conditions (target vs. non-target) decoding results and the behavioral reaction time and accuracy, respectively. (B), Right and left graphs indicate the correlations calculated between category decoding results and the behavioral reaction times and accuracy, respectively. The circles indicate the time points at which the corresponding correlations (FDR-corrected Pearson linear correlation with p < 0.05) were significantly positive or negative.

## Discussion

A growing set of studies have investigated the spatiotemporal dynamics of object recognition in the human brain (Kaneshiro et al., 2015; Karimi-Rozubahani et al., 2017a; Hebart et al., 2018; Cichy et al., 2014; Liu et al. 2009). However, they did not investigate the flow of brain’s top-down attentional effect and its impact on object processing. This work has extended previous results by evaluating the effect of top-down attentional signals in the processing of object categories. For that purpose, first we evaluated the target-related effects (i.e. difference between the target and non-target categories) in the patterns of ERP signals and found target-related modulations in ERPs respectively after the onset of stimulus presentation (in parietal and prefrontal areas) as well as prior to its onset (only at prefrontal area; Fig. 2). By assessing the impacts of target-related effects on the decoding of categories, we observed negligible differences at later time windows (after +300 ms, Fig. 3A) and an earlier appearance of target categories compared to non-target categories (Fig. 3B). Decoding results showed that target-related effects could be decoded from the brain signals of almost all brain areas with higher amplitudes at anterior compared to posterior areas (Figs. 3D and 4). Most interestingly, we saw that target-related preparatory effects appeared in prefrontal areas even prior to the onset of the stimulus (Fig. 4). We observed that, while the target-related signals were sent from the anterior to posterior brain areas during the presentation of the stimulus (Figs. 5 and 6), the categorical signals were sent to the posterior brain areas from 80 ms and forward to anterior areas from 300 ms post-stimulus (Fig. 7).

### Spatiotemporal dynamics of category- and target-related information processing in the brain

Our ERP patterns (Figs. 2), especially on frontal and prefrontal electrodes, repeated the ERP results previously reported for a target detection (fast presentation) go/no-go task (see figure 2 of Thorpe et al., 1996), with dominant N-P-N-P sequence of components (in the first 300 ms post-stimulus onset) and the initiation of target/non-target effects on anterior brain areas. Specifically, using univariate ERP analysis in an animal detection task in which the target did not vary across trials, the mentioned study (Thorpe et al., 1996) suggested that the presence of animal in a naturalistic image could be observed in brain signals as early as 150 ms post-stimulus. Moreover, they showed a possible target-related effect covering the frontal brain areas initiating at around the same time. However, they did not evaluate whether the difference in ERPs were really reflecting categorical/target-related information. They also overlooked the fact that the target-related signals on frontal brain areas could possibly carry task information to posterior areas. Current study, while providing an early target-related ERP effect in prefrontal followed by parietal areas (Fig. 2), extended previous results using MVPA and Granger causality, and showed that the prefrontal effects contained target-related information (rather than only reflecting signal difference) which was sent to posterior areas possibly to facilitate object recognition. Moreover, as our observed effects were specific to PFC (and absent in frontal areas) and first appeared in the pre-stimulus window, it is unlikely that the observed effects were motor-preparation signals; this is not totally true about the mentioned work (Thorpe et al., 1996).

Our results compares with a recent target (object) detection study, in which LFPs were recorded from different areas of the human brain (Bansal et al., 2014). That study, which used short stimulus presentation time in target-based trials which included 50 stimuli, showed target-related effects appearing after 250 ms post-stimulus which mainly reflected as an increase in Gamma band power. While our study suggests that the target-related information processing was initiated in prefrontal, they found the inferotemporal, supramarginal inferior temporal and fusiform gyri to be the earliest and main locations for the encoding of target-related information (Bansal et al., 2014) as compared to inferior frontal and orbital gyri which were also covered in their study (see figure 9 in Bansal et al., 2014). This discrepancy can be explained in light of several considerations. First, they had much fewer electrodes in prefrontal areas, especially on the dorsal PFC, compared to temporal areas (see figure 7 in Bansal et al., 2014), which might have caused missing the early prefrontal effects observed here by excluding some critical areas. Moreover, as a result of having only a few electrodes on the dorsal PFC, the decoding results were probably noisier on frontal areas compared to occipitotemporal areas, causing less amplified effect on frontal areas. Second, we used a longer presentation time compared to the mentioned study; it has been previously suggested that the difference in presentation time can affect the anterior/posterior dynamics of information flow in the human brain (compare (Bar et al., 2006; Karimi-Rouzbahani, 2018) and (Goddard et al., 2016).

While several previous studies have suggested the initiation of task-relevant (i.e. attentional/predictive) activity in higher-order cognitive areas (e.g. PFC) which were sent to lower-level sensory processing areas of the brain (Bar et al., 2006; Summerfield et al., 2006; Gregoriou et al., 2009), they failed to provide quantitative evidence for the movement of task-relevant ‘information’ across the mentioned areas. One of these studies observed a shift of signal power between the two areas and the authors claimed that the categorical signals moved from orbitofrontal cortex to the fusiform gyri (Bar et al., 2006). Results of another study suggested that the causal observation of signal activity across frontal and the LOC area accounted for top-down predictive signals across the two areas (Summerfield et al., 2006). A monkey recording study suggested that shifted gamma-band (8-13 ms) synchronization between FEF and V4 provided a substrate for top-down attentional feedback (Gregoriou et al., 2009). As you probably agree, these observations do not provide enough justification for the movement of information, as the mentioned signals could have reflected an epiphenomenal effect which did not necessarily contain any task-related information. The recently proposed modified version of Granger causality combined with MVPA, which we used here, allowed us to follow the movement of the desired (target-or category-related) information in the brain.

Several of our findings directly compare with a recent study which investigated the spatiotemporal dynamics of object- and task-related information processing in the human brain (Hebart et al., 2018). This includes that we both observed a late (after 300 ms) appearance of task-related information in the human brain (Fig. 3A) which only modulated object representations to a limited degree. We both also observed a gradually increasing task- and category-related information respectively along the dorsal and ventral visual processing hierarchies (Fig. 4). As was previously observed (Vaziri-Pashkam et al., 2017), the task- and category-related information were dominant in the frontoparietal and occipitotemporal cortices, respectively. The current study, extended these results by investigating the flow of task- and category-related information in the brain and showed that the processing of target-related information was initiated on prefrontal areas and the information was sent to posterior brain areas probably for facilitated decision-making and recognition (Bar et al., 2006; Summerfield et al., 2006).

The results of current study, extended two previous EEG/MEG studies, which investigated the flow of categorical (Goddard et al., 2016) as well as category-orthogonal variations (Karimi-Rouzbahani, 2018) across peri-occipital and peri-frontal areas. The MEG study observed the feedforward flow of information from peri-occipital to peri-frontal areas during the stimulus presentation, which lasted for 500 ms, followed by feedback flows in the post-stimulus offset time, and suggested that the feedforward flow, which was unexpected based on previous studies (Bar et al., 2006; Bar et al., 2001; Kveraga et al., 2007), was imposed by the dominance of stimulus-related information input. Our previous EEG study, which used a similar methodology and a shorter presentation time for the stimulus, did not rule out the impact of stimulus presentation on the direction of information flow in the brain (Karimi-Rozubahani, 2018). Showing the feedback of category and target-related information during the stimulus presentation time (Figs. 5 and 7), the current study suggested that, while the stimulus may play role, the direction of information flow could be majorly influenced by the internal task.

### Predictive coding of target versus non-target categories

A post-hoc analysis showed that three and five of the subjects were shown more non-target and target stimuli as their final stimuli (i.e. stimuli 9-12) in trials, respectively (Supplementary Fig. S4A). This unbalanced distribution of target and non-target stimuli within trials, although non-significant (Wilcoxon’s signed rank test, P > 0.067), has probably given these subjects some guesses about nature of the coming stimuli in the same trial (whether they were target or non-target). Based on this information, the pre-stimulus difference in ERP activities (Fig. 2) and decoding (Figs. 3D and 4) of prefrontal areas could have produced some preparatory/modulatory signal sent to visual areas to facilitate the processing/ignoring of the coming stimuli respectively in subjects with higher/lower number of late target stimuli in the trials. This is supported by the ERP signals shown in Supplementary Fig. S4C, which revealed a bias in the pre-stimulus ERP signals for both groups of subjects while showing no pre-stimulus difference in ERPs for the two remaining subjects which were shown equal number of target and non-target final stimuli within trials (Supplementary Fig. S4B).

But what is the underlying mechanisms of prediction? Prior information can direct the brain’s processing towards the subject’s target features/categories when the brain receives multiple inputs (Wald and Wolfowitz, 1949). Accordingly, it is suggested that the brain generates prediction signals which, by weighting the perceptual alternatives, help in predicting the coming stimuli (Friston, 2003). Specifically, the predictive accounts of decision making suggest that, the brain compares the received evidence with an internal ‘template’ and produces a ‘matched’ signal if the matching exceeds a certain threshold (Swets, 1964). Therefore, the neural decision-making architectures, many of which are found across the frontoparietal network, need to access the template information for evaluating the matching of the input stimulus against it. Accordingly, we showed that the target-related information could be decoded from prefrontal areas even before the stimuli appeared (Fig. 4). More importantly, we showed that this information was sent to the posterior (mainly parietal) areas of the brain probably to take part in decision making (Figs. 5 and 6). Moreover, the category-related information was most dominantly sent to occipital areas, probably to enhance the quality of the input stimuli’s representations (Jackson et al., 2017). In conclusion, we showed that the prefrontal areas participate in generating the mentioned template which is used for predictive coding of the input stimulus in the brain.

### Neural correlate for behavioral results

We had hypothesized that category-related decoding, which represented externally driven information processing in the brain, could produce significantly higher decoding values compared to the target-related results, which reflected the internally induced information. However, it was shown that target-related information provided decoding values not only comparable, but at some windows also higher than the category-related information (Fig. 4). In fact, the target-related information, which is task-relevant, seem to have dominated the brain processing. It seems that, what the subjects are concentrating on is not the classification of object categories, but rather, detecting the target categories among non-target categories. This is probably the reason why, rather than with the decoding of categories, the behavioral results mainly correlated with the decoding of target-related information especially at final time windows (Fig. 8). In other words, categorization played a peripheral rather than a central role in the category recognition task of this study.

### Future directions

While our results shed some light on the role of prefrontal cortices in the initiation of task-relevant information processing in object recognition, a fusion of EEG/MEG and fMRI data as recently done on a similar question (Hebart et al., 2018), equipped with the recently developed version of RSA-Granger causality analysis (Goddard et al., 2016), may add precision to the exact sources of this information processing in the brain.

It was shown here that target-related information processing was initiated in PFC and was sent to frontoparietal network. However, as opposed to our expectation, we only saw an ignorable late difference between the category decoding across the target and non-target conditions (Fig. 3A). Therefore, it has remained unknown whether (if at all) this information could contribute to category representations in visual areas. Using transcranial magnetic stimulation (TMS) applied to the frontal brain areas in a chosen set of trials, by deactivating these areas, could provide the opportunity of studying the impact target-related information sent to those areas.

Our results, while extending previous findings on the spatiotemporal dynamics of task- and category-related information processing in the brain (Hebart et al., 2018), provided new insights into the role of prefrontal areas in object recognition.

## Acknowledgements

We would like to thank Ali Mazdarani for providing us with the cluster system which we used for our analyses.

## References

Azouz, R. and Gray, C.M., 1999. Cellular mechanisms contributing to response variability of cortical neurons in vivo. Journal of Neuroscience, 19(6), pp. 2209–2223.

Bansal, A.K., Madhavan, R., Agam, Y., Golby, A., Madsen, J.R. and Kreiman, G., 2014. Neural dynamics underlying target detection in the human brain. Journal of Neuroscience, 34(8), pp. 3042–3055.

Bar, M., Tootell, R.B., Schacter, D.L., Greve, D.N., Fischl, B., Mendola, J.D., Rosen, B.R. and Dale, A.M., 2001. Cortical mechanisms specific to explicit visual object recognition. Neuron, 29(2), pp. 529–535.

Bar, M., Kassam, K.S., Ghuman, A.S., Boshyan, J., Schmid, A.M., Dale, A.M., Hämäläinen, M.S., Marinkovic, K., Schacter, D.L., Rosen, B.R. and Halgren, E., 2006. Top-down facilitation of visual recognition. Proceedings of the national academy of sciences, 103(2), pp. 449–454.

Battistoni, E., Stein, T. and Peelen, M.V., 2017. Preparatory attention in visual cortex. Annals of the New York Academy of Sciences, 1396(1), pp. 92–107.

Brainard, D.H. and Vision, S., 1997. The psychophysics toolbox. Spatial vision, 10, pp. 433–436.

Cichy, R.M., Pantazis, D. and Oliva, A., 2014. Resolving human object recognition in space and time. Nature neuroscience, 17(3), p. 455.

Chang, C.C. and Lin, C.J., 2011. LIBSVM: a library for support vector machines. ACM transactions on intelligent systems and technology (TIST), 2(3), p. 27.

Daliri, M.R., Taghizadeh, M. and Niksirat, K.S., 2013. EEG signature of object categorization from event-related potentials. Journal of medical signals and sensors, 3 (1), p. 37.

Davis, N.J., Tomlinson, S.P. and Morgan, H.M., 2012. The role of beta-frequency neural oscillations in motor control. Journal of Neuroscience, 32(2), pp. 403–404.

Delorme, A. and Makeig, S., 2004. EEGLAB: an open source toolbox for analysis of single-trial EEG dynamics including independent component analysis. Journal of neuroscience methods, 134(1), pp. 9–21.

DiCarlo, J.J., Zoccolan, D. and Rust, N.C., 2012. How does the brain solve visual object recognition?. Neuron, 73(3), pp. 415–434.

Duncan, J. and Owen, A.M., 2000. Common regions of the human frontal lobe recruited by diverse cognitive demands. Trends in neurosciences, 23(10), pp. 475–483.

Friston, K., 2003. Learning and inference in the brain. Neural Networks, 16(9), pp. 1325–1352.

Goddard, E., Carlson, T.A., Dermody, N. and Woolgar, A., 2016. Representational dynamics of object recognition: Feedforward and feedback information flows. Neuroimage, 128, pp. 385–397.

Granger, C.W., 1969. Investigating causal relations by econometric models and cross-spectral methods. Econometrica: Journal of the Econometric Society, pp. 424–438.

Gregoriou, G.G., Gotts, S.J., Zhou, H. and Desimone, R., 2009. Long-range neural coupling through synchronization with attention. Progress in brain research, 176, pp. 35–45.

Harel, A., Ullman, S., Harari, D. and Bentin, S., 2011. Basic-level categorization of intermediate complexity fragments reveals top-down effects of expertise in visual perception. Journal of vision, 11(8), pp. 18–18.

Hebart, M.N., Bankson, B.B., Harel, A., Baker, C.I. and Cichy, R.M., 2018. The representational dynamics of task and object processing in humans. Elife, 7, p. e32816.

Hong, H., Yamins, D.L., Majaj, N.J. and DiCarlo, J.J., 2016. Explicit information for category-orthogonal object properties increases along the ventral stream. Nature neuroscience, 19(4), p. 613.

Isik, L., Meyers, E.M., Leibo, J.Z. and Poggio, T., 2013. The dynamics of invariant object recognition in the human visual system. Journal of neurophysiology, 111(1), pp. 91–102.

Jackson, J., Rich, A.N., Williams, M.A. and Woolgar, A., 2017. Feature-selective attention in frontoparietal cortex: multivoxel codes adjust to prioritize task-relevant information. Journal of cognitive neuroscience, 29(2), pp. 310–321.

Kaneshiro, B., Guimaraes, M.P., Kim, H.S., Norcia, A.M. and Suppes, P., 2015. A representational similarity analysis of the dynamics of object processing using single-trial EEG classification. Plos one, 10(8), p. e0135697.

Karimi-Rouzbahani, H., 2018. Three-stage processing of category and variation information by entangled interactive mechanisms of peri-occipital and peri-frontal cortices. Scientific Reports, 8, 12213.

Karimi-Rouzbahani, H., Bagheri, N. and Ebrahimpour, R., 2017a. Average activity, but not variability, is the dominant factor in the representation of object categories in the brain. Neuroscience, 346, pp. 14–28.

Karimi-Rouzbahani, H., Bagheri, N. and Ebrahimpour, R., 2017b. Hard-wired feed-forward visual mechanisms of the brain compensate for affine variations in object recognition. Neuroscience, 349, pp. 48–63.

Karimi-Rouzbahani, H., Bagheri, N. and Ebrahimpour, R., 2017c. Invariant object recognition is a personalized selection of invariant features in humans, not simply explained by hierarchical feed-forward vision models. Scientific reports, 7(1), p. 14402.

Karimi-Rouzbahani, H., Ebrahimpour, R. and Bagheri, N., 2016. Quantitative evaluation of human ventral visual stream in invariant object recognition: Human behavioral experiments and brain-plausible computational model simulations. Machine Vision and Image Processing, 3, pp. 59–72.

Kelly, S.P. and O’Connell, R.G., 2013. Internal and external influences on the rate of sensory evidence accumulation in the human brain. Journal of Neuroscience, 33(50), pp. 19434–19441.

Kiani, R., Esteky, H., Mirpour, K. and Tanaka, K., 2007. Object category structure in response patterns of neuronal population in monkey inferior temporal cortex. Journal of neurophysiology, 97(6), pp. 4296–4309.

Klimesch, W., 2012. Alpha-band oscillations, attention, and controlled access to stored information. Trends in cognitive sciences, 16(12), pp. 606–617.

Kravitz, D.J., Kriegeskorte, N. and Baker, C.I., 2010. High-level visual object representations are constrained by position. Cerebral Cortex, 20(12), pp. 2916–2925.

Kveraga, K., Boshyan, J. and Bar, M., 2007. Magnocellular projections as the trigger of top-down facilitation in recognition. Journal of Neuroscience, 27(48), pp. 13232–13240.

Lamme, V.A. and Roelfsema, P.R., 2000. The distinct modes of vision offered by feedforward and recurrent processing. Trends in neurosciences, 23(11), pp. 571–579.

Liu, H., Agam, Y., Madsen, J.R. and Kreiman, G., 2009. Timing, timing, timing: fast decoding of object information from intracranial field potentials in human visual cortex. Neuron, 62(2), pp. 281–290.

Makeig, S., 1993. Auditory event-related dynamics of the EEG spectrum and effects of exposure to tones. Electroencephalography and clinical neurophysiology, 86(4), pp. 283–293.

Manahova, M.E., Mostert, P., Kok, P., Schoffelen, J.M. and de Lange, F.P., 2018. Stimulus familiarity and expectation jointly modulate neural activity in the visual ventral stream. Journal of cognitive neuroscience, pp. 1–12.

Meijs, E.L., Slagter, H.A., de Lange, F.P. and van Gaal, S., 2018. Dynamic interactions between top-down expectations and conscious awareness. Journal of Neuroscience, pp. 1952–17.

Meyers, E., 2013. The neural decoding toolbox. Frontiers in neuroinformatics, 7, p. 8.

Milton, A. and Pleydell-Pearce, C.W., 2016. The phase of pre-stimulus alpha oscillations influences the visual perception of stimulus timing. Neuroimage, 133, pp. 53–61.

Mognon, A., Jovicich, J., Bruzzone, L. and Buiatti, M., 2011. ADJUST: An automatic EEG artifact detector based on the joint use of spatial and temporal features. Psychophysiology, 48(2), pp. 229–240.

Moran, J. and Desimone, R., 1985. Selective attention gates visual processing in the extrastriate cortex. Science, 229(4715), pp. 782–784.

Nácher, V., Ledberg, A., Deco, G. and Romo, R., 2013. Coherent delta-band oscillations between cortical areas correlate with decision making. Proceedings of the National Academy of Sciences, p. 201314681.

Paneri, S. and Gregoriou, G.G., 2017. Top-down control of visual attention by the prefrontal cortex. functional specialization and long-range interactions. Frontiers in neuroscience, 11, p. 545.

Puri, A.M., Wojciulik, E. and Ranganath, C., 2009. Category expectation modulates baseline and stimulus-evoked activity in human inferotemporal cortex. Brain research, 1301, pp. 89–99.

Rohenkohl, G. and Nobre, A.C., 2011. Alpha oscillations related to anticipatory attention follow temporal expectations. Journal of Neuroscience, 31(40), pp. 14076–14084.

Schwarzlose, R.F., Swisher, J.D., Dang, S. and Kanwisher, N., 2008. The distribution of category and location information across object-selective regions in human visual cortex. Proceedings of the National Academy of Sciences, 105(11), pp. 4447–4452.

Soon, C.S., Namburi, P. and Chee, M.W., 2013. Preparatory patterns of neural activity predict visual category search speed. Neuroimage, 66, pp. 215–222.

Spyropoulos, G., Bosman, C.A. and Fries, P., 2018. A theta rhythm in macaque visual cortex and its attentional modulation. Proceedings of the National Academy of Sciences, p. 201719433.

Steinmetz, N.A. and Moore, T., 2009. Changes in the response rate and response variability of area V4 neurons during the preparation of saccadic eye movements. Journal of neurophysiology, 103(3), pp. 1171–1178.

Stokes, M., Thompson, R., Nobre, A.C. and Duncan, J., 2009. Shape-specific preparatory activity mediates attention to targets in human visual cortex. Proceedings of the National Academy of Sciences, 106(46), pp. 19569–19574.

Storey, J.D., 2002. A direct approach to false discovery rates. Journal of the Royal Statistical Society: Series B (Statistical Methodology), 64(3), pp. 479–498.

Summerfield, C. and Egner, T., 2009. Expectation (and attention) in visual cognition. Trends in cognitive sciences, 13(9), pp. 403–409.

Summerfield, C., Egner, T., Greene, M., Koechlin, E., Mangels, J. and Hirsch, J., 2006. Predictive codes for forthcoming perception in the frontal cortex. Science, 314(5803), pp. 1311–1314.

Swets, J. and Green, D., 1964. in Signal Detection and Recognition, Eds. (Wiley, New York), pp. 164–171.

Tanner, D., Morgan-Short, K. and Luck, S.J., 2015. How inappropriate high-pass filters can produce artifactual effects and incorrect conclusions in ERP studies of language and cognition. Psychophysiology, 52(8), pp. 997–1009.

Thorpe, S., Fize, D. and Marlot, C., 1996. Speed of processing in the human visual system. Nature, 381(6582), p. 520.

Thorpe, S.J., Rolls, E.T. and Maddison, S., 1983. The orbitofrontal cortex: neuronal activity in the behaving monkey. Experimental Brain Research, 49(1), pp. 93–115.

VanRullen, R., 2007. The power of the feed-forward sweep. Advances in Cognitive Psychology, 3(1-2), p. 167.

VanRullen, R. and Thorpe, S.J., 2001. The time course of visual processing: from early perception to decision-making. Journal of cognitive neuroscience, 13(4), pp. 454–461.

Vaziri-Pashkam, M. and Xu, Y., 2017. Goal-directed visual processing differentially impacts human ventral and dorsal visual representations. Journal of Neuroscience, pp. 3392–16.

Wald, A. and Wolfowitz, J., 1950. Bayes solutions of sequential decision problems. The Annals of Mathematical Statistics, pp. 82–99.

Widmann, A. and Schröger, E., 2012. Filter effects and filter artifacts in the analysis of electrophysiological data. Frontiers in psychology, 3, p. 233.

Woolgar, A., Afshar, S., Williams, M.A. and Rich, A.N., 2015a. Flexible coding of task rules in frontoparietal cortex: an adaptive system for flexible cognitive control. Journal of cognitive neuroscience, 27(10), pp. 1895–1911.

Woolgar, A., Williams, M.A. and Rich, A.N., 2015b. Attention enhances multi-voxel representation of novel objects in frontal, parietal and visual cortices. Neuroimage, 109, pp. 429–437.

